# Taxonomic Composition and Predicted Functional Potential of a Commercial Microbiome-Based Fertilizer Additive and Agricultural Soils in Eastern Paraguay

**DOI:** 10.64898/2026.06.03.729874

**Authors:** Walter J. Sandoval-Espinola

**Affiliations:** Facultad de Ciencias Exactas y Naturales, Departamento de Biotecnología, Laboratorio de Biotecnología Microbiana, Universidad Nacional de Asunción, San Lorenzo, Paraguay; MicroBios, Montevideo, Uruguay

**Keywords:** soil microbiome, fertilizers, clime-smart agriculture

## Abstract

Anthropogenic soil degradation is a major challenge for sustainable food production, particularly in tropical agricultural systems where excessive fertilizer use contributes to soil deterioration and greenhouse gas emissions. Microbiome-based agricultural technologies offer a potential strategy to improve fertilizer-use efficiency while maintaining crop productivity. Here, we characterized the taxonomic composition and predicted functional potential of a commercial microbiome-based fertilizer additive (humus) deployed across more than 1.4 million hectares in Paraguay and Uruguay, and compared it with root-associated microbiomes. In parallel, we evaluated agricultural and forest soil microbiomes from eastern Paraguay. Microbial communities were analyzed using 16S rRNA gene sequencing and PICRUSt2-based functional prediction. The humus microbiome displayed enrichment of pathways associated with degradation of organic compounds, nutrient cycling, and plant-growth-promoting activities. Furthermore, humus and root-associated microbiomes shared over 350 predicted microbial pathways, indicating substantial functional overlap despite differences in specific bacterial taxa, and suggesting that the consortium may function as a rhizosphere-like microbial community capable of providing functions commonly associated with plant-associated microbiomes. In agricultural soils, significant taxonomic differences were observed between high- and low-productivity fields, whereas predicted functional profiles remained largely conserved, consistent with functional redundancy within soil microbial communities. Productive soils were enriched in the superpathway of demethylmenaquinol-6 biosynthesis II, a microbial vitamin K2-related pathway involved in respiratory metabolism. Together, these findings provide the first detailed taxonomic and predicted functional characterization of a large-scale commercial microbiome-based fertilizer additive and establish a baseline for understanding microbial diversity and functional potential across productive agricultural soils in Paraguay.

**Importance:** Soil degradation and inefficient fertilizer use are major constraints to sustainable agriculture, particularly in tropical systems where nutrient losses and greenhouse gas emissions are high. Microbiome-based agricultural inputs are increasingly proposed as tools to enhance soil functioning and improve nutrient cycling efficiency, yet their ecological characteristics and functional potential remain poorly understood. This study provides the first detailed taxonomic and predicted functional characterization of a large-scale commercial microbiome-based fertilizer additive deployed in South American agriculture, and places it in the context of native forest and agricultural soil microbiomes.

By integrating 16S rRNA gene sequencing with predictive functional profiling, this work reveals substantial functional overlap between the microbial consortium and plant-associated microbiomes, suggesting ecological convergence toward rhizosphere-like functions. In addition, the identification of conserved functional profiles across soils with contrasting productivity highlights the potential role of functional redundancy in maintaining ecosystem processes under different management regimes. Together, these findings provide a foundational framework for understanding microbiome-based agricultural inputs and soil microbial functional stability in subtropical agroecosystems, specifically Paraguay, contributing to the development of more sustainable agricultural practices.

## Introduction

One of the greatest challenges to sustainable food production is soil degradation, which involves the loss of essential biological, physical, and chemical properties^1^. These changes also profoundly affect soil microbial community composition, leading to lower crop yields^2–5^. Notably, soil degradation is most prevalent in tropical regions, where many developing countries are located.

Natural and anthropogenic factors drive soil degradation, including compaction, erosion, salinization, leaching, proliferation of soil-borne pathogens, loss of soil macro- and micro-biodiversity, and disruption of nutrient cycling. These changes lead to a decline in soil productivity, affecting food security^2^. Ironically, one of the key drivers of soil degradation is the poor management of synthetic fertilizers^3^. While their deployment has increased yields, overuse beyond recommended levels leads to compaction and salinization^6^. This, in turn, compels producers to further increase dosages, potentially turning this important practice into a vicious cycle. Each year, over 90 million metric tons of nitrogenous fertilizers are added to the soil. Unfortunately, up to 70% is lost via nitrate leaching, ammonia volatilization, and N_2_O gas emissions^7–9^. Moreover, this gas represents the third most prevalent anthropogenic greenhouse gas (GHG), with up to 300 CO_2_ equivalents (CO2eq)^10^ in terms of its heat-trapping capacity. Therefore, the increase in nitrogenous fertilizers use under this suboptimal condition, can aggravate soil degradation and climate change^3,11,12^. Thus, approaches for optimizing the efficiency of traditional fertilizers, while reducing GHG emissions, constitute a powerful strategy for sustainable food production, while preserving the soil and minimizing climate change.

Although there is not a regulatory definition for regenerative agriculture, its practice points towards sustainable food production and soil preservation^13,14^. These approaches include the elimination of tillage, the use of cover crops, or by increasing biodiversity via mix farming. Interestingly, at the core of these practices lies soil microbiology. Here, the soil microbiome plays a key role in nutrient cycling and in establishing a synergistic relationship with plants. These practices, while preserving and enhancing the soil’s characteristics, allow for improved crop yields^15–17^. Therefore, strategies that harness the metabolic capabilities of the soil microbiome can be important tools towards sustainable food production and soil preservation^18–21^.

A previously developed microbiome-based fertilizer additive (humus), originally developed to resemble characteristics of healthy root-associated microbiomes, increases fertilizer bioavailability, resulting in higher crop yields^22^. Specifically, previous field evaluations reported that replacing 30% of conventional nitrogen-phosphorous-potassium (NPK) fertilizer with this biological additive was associated with approximately 12% higher crop yields. The humus has been commercially applied (Tiroleo’s M.o. Humus) in a combined area of over 1.4 million hectares over the past six years in South America, including Paraguay and Uruguay^23^.

While the agronomic benefits of this microbiome-based fertilizer additive have been previously reported, its microbial composition and predicted functional potential have not been comprehensively characterized. Moreover, to our knowledge, only one previous report in Paraguay evaluated the effects of a biostimulant on soil microbiome^25^. Thus, despite the agricultural importance of Paraguay^24^, information regarding the taxonomic and functional structure of its soil microbiomes remains limited. Understanding how microbial communities vary across soils with different productivity levels may help identify microbial signatures^26,27^ associated with ecosystem functioning and crop performance.

Here, we characterized the microbial composition and predicted functional potential of the humus using 16S rRNA gene sequencing and PICRUSt2^28^. We further compared the humus microbiome with root-associated communities and evaluated agricultural and forest soils from eastern Paraguay. Specifically, we sought to identify microbial taxa and predicted functions associated with the fertilizer additive, determine its overlap with root microbiomes, and assess taxonomic and functional patterns across soils differing in productivity.

In summary, this work provides insights into microbial functions potentially associated with the improved fertilizer-use efficiency observed following humus application, and the microbial ecology of productive soils in an important agricultural region of South America.

## Results

### The microbiome-based fertilizer additive composition and functional potential

A microbiome-based fertilizer additive was previously developed and commercially applied (Tiroleo’s M.o. Humus) in a combined area of over 1.5 million hectares across Paraguay and Uruguay^22,23^. Its use allows for farmers to decrease their chemical fertilizer input by 30% while increasing their crops yield by approximately 12% on average. For short, this additive will be called “humus” in this text. It’s generation involves a two-stage fermentation process ^22^. The first involves the inoculum generation, and the second consists of a static solid-state fermentation using plant biomass as substrate. This stage is performed in 3 m deep boxes filled with the substrate, in a combined volume of approximately 45 m^3^. The humus’ microbial composition was previously mentioned, however, only at the phylum level. In that previous report, rhizosphere microbiomes from two well-grown succulent plants (Crassulaceae family) had also been analyzed. In this work, they are annotated as samples number 13 and 14. For the humus microbiome analysis, three samples were obtained at the end of the solid-state fermentation phase, which takes three weeks, covering the box’s bottom, middle, and top (surface) parts. These corresponded to samples 6, 5, and 4, respectively. The reanalyzed data was tabulated in terms of type, either root or humus, and depth, either simply root or, for the humus, surface, middle, or bottom. For this work, we performed a deeper evaluation of the humus taxonomical and functional potential through PICRUSt2^28^ analysis.

Microbiome analysis through 16S rRNA V3-V4 sequencing, with an average depth of 35,497 reads per sample, revealed slight non-significant differences between each humus sample (Figure 1-A). At the phylum level, the five most abundant groups within the surface sample were Proteobacteria (31%), Chloroflexi (17%), Actinobacteriota (9%), Gemmatimonadota (5%), and Planctomycetota (5%). The middle’s five most abundant were Actinobacteriota (41%), Bacteroidota (36%), Firmicutes (15%), Proteobacteria (3%), and Aquificota (3%). Finally, the bottom sample contained Firmicutes (52%), Actinobacteriota (41%), Proteobacteria (3%), Myxococcota (1%), and Bacteroidota (0.6%) as the top 5 most abundant. At the genus level (Figure 1-B), the top sample contained mostly *Acidibacter, Heliangium*, and *Verrucosispora*. The middle contained mostly *Chitinophaga, Glycomyces*, and *Streptomyces*. Whereas the bottom contained mostly *Streptomyces, Effusibacillus*, and *Paenibacillus*. In terms of beta diversity (Figure 1-C), there was no significant differences between the root and humus microbiomes composition according to a permutational multivariate analysis of variance (PERMANOVA) (Bray-Curtis distance; F=1.27, p = 0.2). Similarly, in terms of alpha diversity, while the Shannon index was higher in root compared to the combined humus samples, it was not significant (Kruskal-Wallis rank sum test p = 0.08). When looking at depth (Figure 1-D), the Shannon index was higher in root and surface samples, however, also not significant (Kruskal-Wallis rank sum test p = 0.28). Overall, these analyses show that the humus microbiome can differ slightly in its composition, likely due to the oxygen gradient within the box, and they were not significantly different from the root microbiomes evaluated.

**Figure 1.**
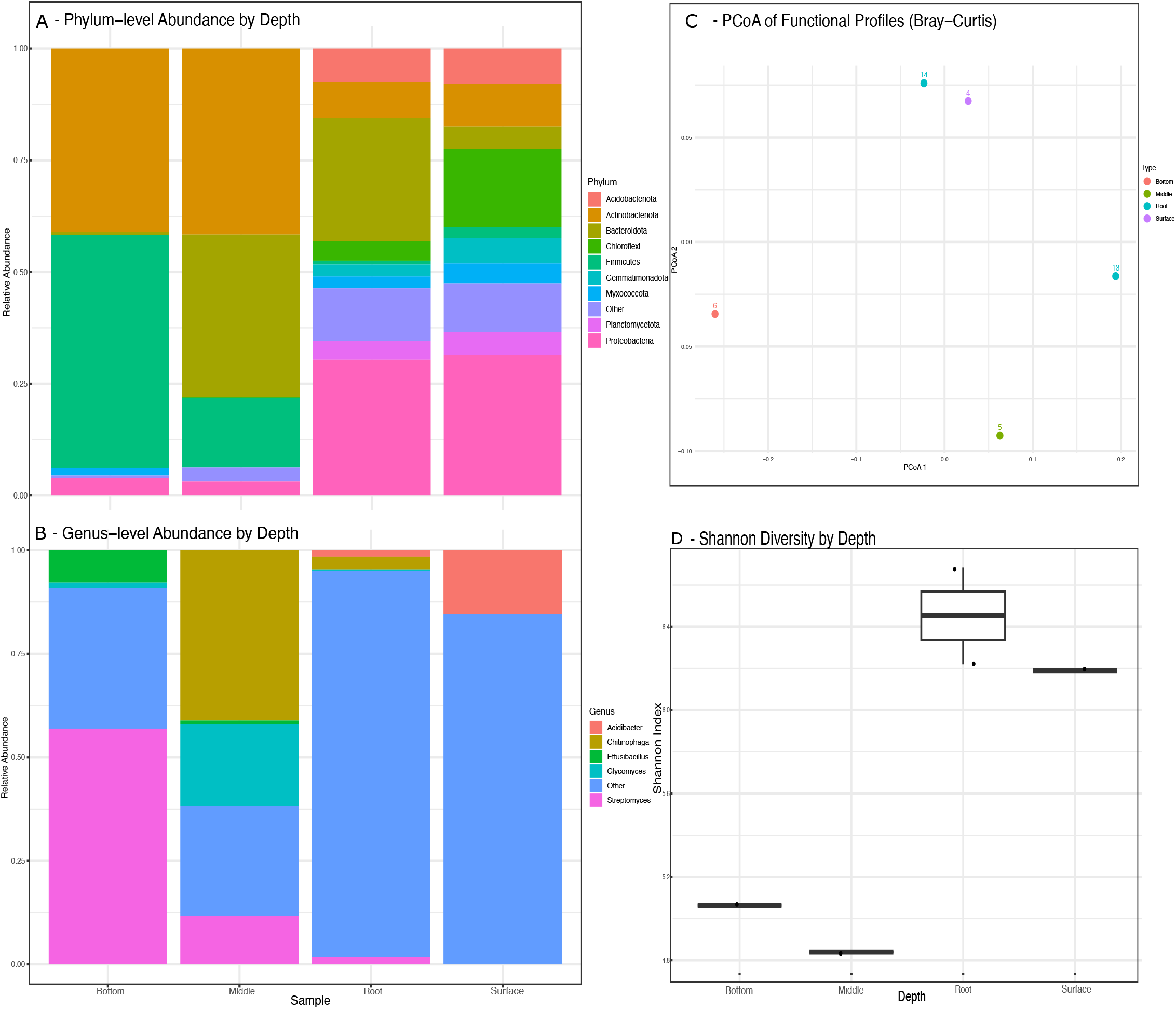
Structure and diversity of humus and root microbiomes. A) relative abundance at phylum and genus across four depth categories: Surface, Middle, Bottom (humus), and roots) (B) Genus-level relative abundances across the same depth categories. C) Beta diversity, Principal Coordinates Analysis (PCoA) of functional profiles based on Bray-Curtis dissimilarity. Samples cluster according to depth Bray-Curtis. D) Shannon diversity indices across depths using box plots.

While no significant differences in overall community composition were observed between groups based on PERMANOVA analysis, an exploratory multivariable association with linear models analysis (MaAsLin2)^29^ (FDR < 0.25), considering root vs humus, revealed an enrichment of genus *Sphingomonas*, env.OPS_17 (Family Sphingobacteriales), and Solirubrobacterales.67.14 within roots. This suggests that differences between groups are driven by these specific microbial lineages rather than broad community-wide shifts.

A PICRUSt2 analysis was then performed to determine the humus and root microbiomes microbial functional potential. Overall, 371, 347, and 350 microbial pathways were identified in the top, middle, and bottom samples, respectively (Supplemental Material 1). The roots microbiomes contained 409 and 401 microbial pathways identified. When combining each group as humus and root microbiomes, 56 pathways were identified as differentially abundant between groups, according to a MaAsLin2 (FDR < 0.25) analysis (Figure 2-A). Specifically, amino acid degradation, aminotransferase, central metabolism, chlorophyll biosynthesis, denitrification, nucleotide, amino acid, and cofactor metabolism, and aromatic compound degradation were enriched in roots, likely reflecting their close interaction with plant hosts and the rhizosphere environment. Alternatively, the humus contained higher abundance of pathways related to glycolysis, central carbon metabolism, TCA cycle and respiratory metabolism, sugar and hexitol degradation, specialized carbon metabolism, and aromatic/complex compounds degradation. These are consistent with organic matter fermentation in the solid-state system (Supplemental Material 2).

**Figure 2.**
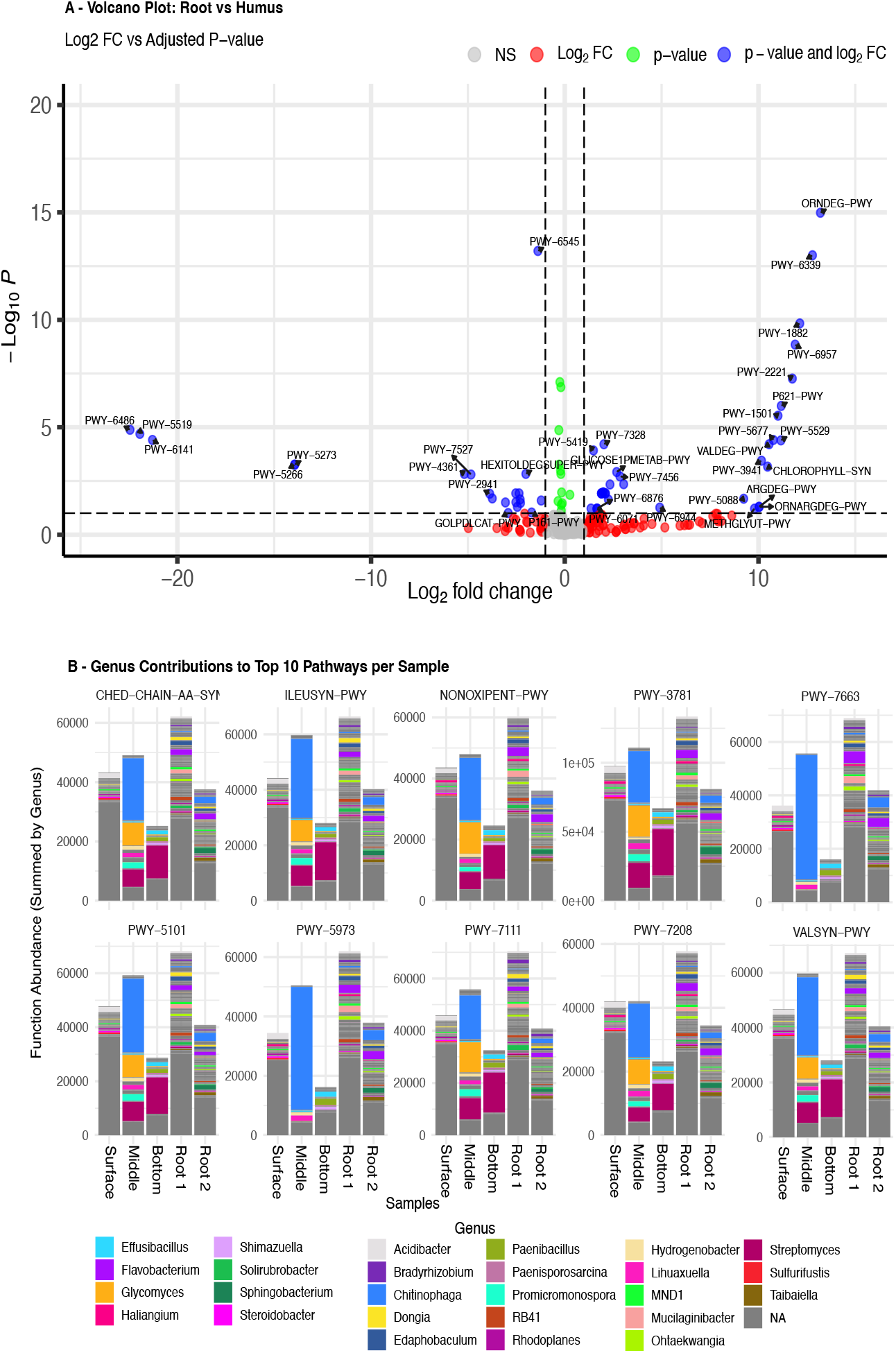
A) Volcano plot showing differential abundance of PICRUSt2 predicted microbial pathways in humus vs root, after MaAsLin2 analysis. B) Top 10 most abundant microbial pathways and their main contributors at the Genus level. BRANCHED-CHAIN-AA-SYN-PWY (branched-chain amino acid biosynthesis), ILEUSYN-PWY (L-isoleucine biosynthesis I), NONOXIPENT-PWY (non-oxidative pentose phosphate pathway), PWY-3781 (aerobic respiration I, cytochrome c), PWY-7663 (anaerobic gondoate biosynthesis), PWY-5101 (L-isoleucine biosynthesis II), PWY-5973 (cis-vaccenate biosynthesis), PWY-7111 (pyruvate fermentation to isobutanol), PWY-7208 (pyrimidine nucleotide salvage superpathway), and VALSYN-PWY (L-valine biosynthesis).

While the above-mentioned patterns highlight the potential metabolic specialization of microbial communities across these habitats, 350 microbial pathways were common among the root and humus microbiomes, potentially underscoring their similarity. Among these, Figure 2-B shows the top 10 most abundant pathways across samples, along with the genera contributing to each. These pathways included branched chain amino acid synthesis, L-isoleucine biosynthesis I and II, non-oxidative pentose phosphate pathway, aerobic respiration, cis-vaccenate biosynthesis, pyruvate fermentation to iso-butanol (engineered), superpathway of pyrimidine nucleobases salvage, gondoate biosynthesis (anaerobic), and L-valine biosynthesis. At the genus level, the 5 most abundant contributors among these pathways are *Chitinophaga, Glycomyces, Streptomyces, Flavobacterium*, and *Promicromonospora*.

For this evaluation we also considered the humus use as additive for chemical fertilizers. As such, we investigated microbial pathways associated to plant-growth-promoting activities^30–32^ (Figure 3). Among these, we identified pathways related to nitrogen metabolism, indole-3-acetic acid (IAA) auxin biosynthesis, siderophores biosynthesis, phosphate solubilization and use, sulfur metabolism, amino acid synthesis, and photosynthesis along with carbon fixation. Overall, these results suggest that these microbial activities might be involved in the physiological results obtained in the field, whereby the humus usage results in higher crop yields with lower NPK input.

**Figure 3.**
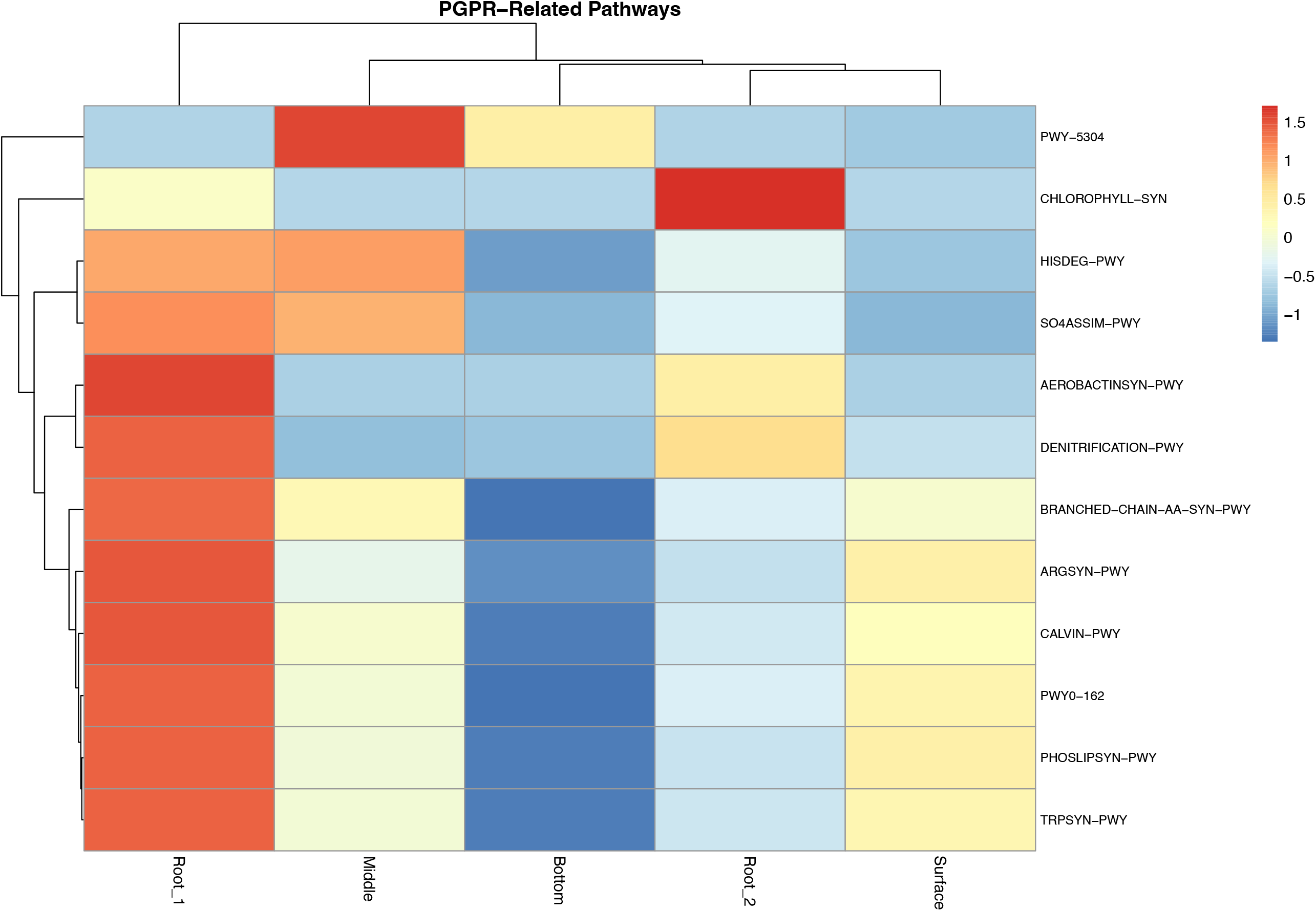
Heatmap showing the relative abundance of identified PGPR-associated metabolic pathways across five samples (Root_13, Root_14, Humus_4 (surface), Humus_5 (middle depth), and Humus_6 (deep), inferred from 16S rRNA gene sequencing data using PICRUSt2. Pathways are hierarchically clustered (dendrograms) based on similarity in predicted functional profiles. The color gradient represents normalized relative abundance values, ranging from −1 (blue, lower abundance) to 1.5 (red, higher abundance), with intermediate levels shown in white/yellow.

Together, these findings provide the first detailed taxonomic and functional characterization of a commercial microbiome-based fertilizer additive currently applied at large agricultural scale in South America.

### Soil microbiome structure and diversity from productive fields of Eastern Paraguay

In addition to expanding the humus’ microbiome composition and predicted microbial functional potential, soil samples obtained from the department of Itapúa in the Eastern Region of Paraguay, were also evaluated. While this work did not include a longitudinal evaluation of the humus application, this section serves as a fundamental contribution to understanding Paraguayan soil microbiome composition. The soil in this region is classified as ultisol, or red clay soil, which is common in tropical and subtropical regions, with acidic pH, and generally known to be highly fertile^33^. Samples were categorized in terms of productivity levels, including “high-productive” and “low-productive”, referring to farmers report regarding their crop yields withing these fields. In addition, we added forest samples obtained from a remanent primary forest in the same region. Overall, this not a comprehensive evaluation of distinct type of soils from Paraguay, but an initial exploratory evaluation from this very productive region.

The soil microbiome analysis was performed through 16S rRNA V3-V4 sequencing, with a mean read count per sample of 25,127 reads, however those with <10,000 reads were excluded from further analysis. Sequencing coverage across samples (Good’s coverage: mean = 0.9998, min = 0.9995) suggested that most microbial diversity was captured. Alpha diversity remained similar among groups (Kruskal–Wallis, p = 0.436) (Figure 4 - A). Alternatively, beta diversity showed that microbial community composition differed across groups (PERMANOVA, R^2^ = 0.168, p = 0.021) (Figure 4 - B), although differences in multivariate dispersion were also detected (PERMDISP, p = 0.019) (Figure 4 - C). This indicates that differences in within-group heterogeneity may contribute to the observed separation. Likewise, pairwise PERMANOVA analyses revealed that the significant compositional shift was primarily driven by differences between high-productive and low-productive soils (R^2^ = 0.105, p = 0.046), whereas comparisons involving Forest soils were not significant (Forest vs high-productive: R^2^ = 0.100, p = 0.165; Forest vs low-productive: R^2^ = 0.296, p = 0.200).

**Figure 4.**
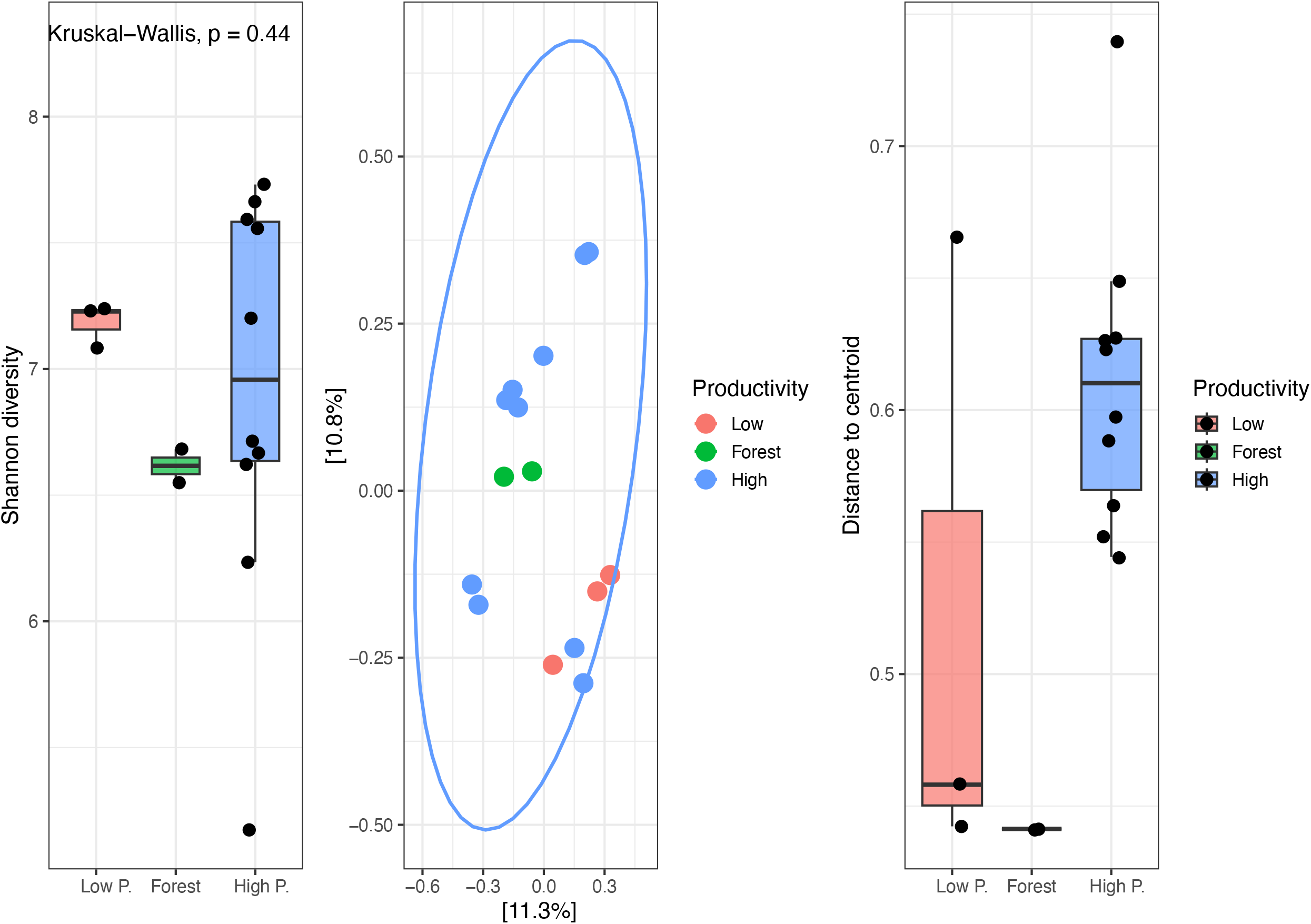
Alpha and beta diversity patterns across soil productivity groups. A) Shannon alpha diversity index across Forest, High-productive, and Low productive soils. No significant differences were detected among groups (Kruskal–Wallis test, p > 0.05). Points represent individual samples and boxplots indicate median and interquartile range. (B) Principal coordinate analysis (PCoA) based on Bray–Curtis dissimilarities showing differences in microbial community composition among productivity groups. Ellipses represent group dispersion. PERMANOVA indicated significant differences in community composition among groups (R^2^ = 0.168, p = 0.021). (C) Distance-to-centroid analysis showing multivariate dispersion within productivity groups. Differences in dispersion were significant according to PERMDISP analysis (p = 0.03), indicating higher community heterogeneity in some productivity categories.

A subsequent MaAsLin2^29^ analysis was performed setting high-productive soils as the reference category (Figure 5). This was selected because pairwise PERMANOVA analyses indicated that the primary differentiation occurred between this and low-productive soils, as mentioned above. Interestingly, despite the lack of broad compositional separation involving Forest soils, MaAsLin2 analysis (FDR < 0.25) identified genus-level associations with soil productivity when high-productive soils were used as the reference category. Forest soils exhibited significant enrichment of *Pseudomonas, Lysobacter, Cellvibrio, Terrabacter, Intrasporangium*, and members of the OM60(NOR5) clade, indicating taxa-specific responses not reflected at the community distance level. Low-productive soils were enriched in taxa including *Bacillus, Paenibacillus, Lysinibacillus, Cohnella, Sphaerobacter, Nitrosomonas, Methylocapsa*, and *Methylovirgula*. Although most taxa were uniquely associated with a single productivity category, *Hyphomicrobium* showed significant associations in both Forest and Low-productive soils relative to high-productive soils. Effect sizes varied across taxa, with the strongest positive associations observed for *Cohnella* in Low-productive soils and *Pseudomonas* in Forest soils.

**Figure 5.**
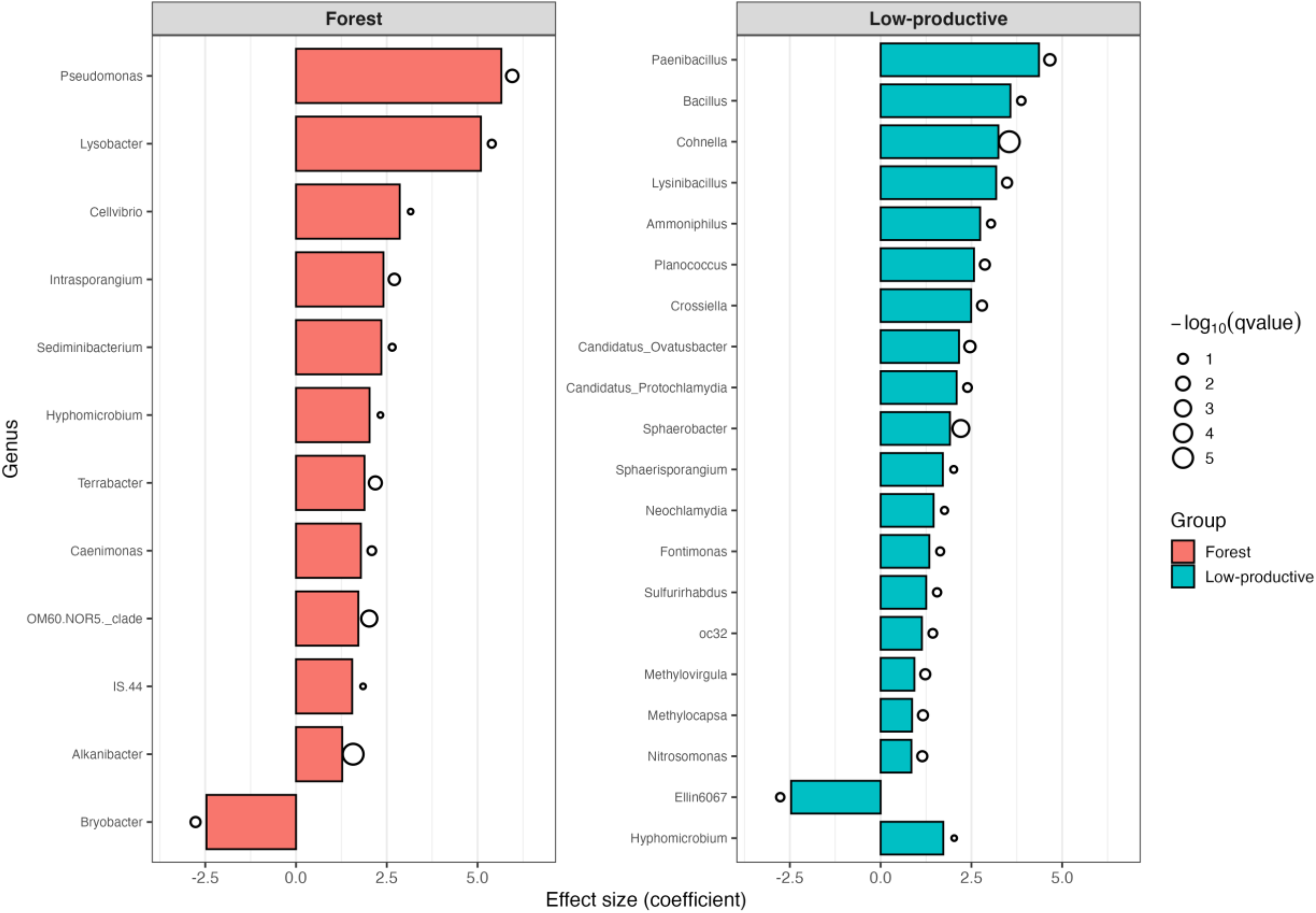
Differentially abundant bacterial genera associated with soil productivity categories identified by MaAsLin2 analysis. Bars represent MaAsLin2 effect sizes (coefficients) relative to the reference group (High-productive soils). Positive coefficients indicate enrichment in either Forest or Low-productive soils compared to High-productive soils, whereas negative coefficients indicate taxa enriched in High-productive soils. Bubble size corresponds to statistical significance expressed as −log10(q-value), where larger bubbles indicate stronger statistical support after false discovery rate correction. Taxa are separated according to comparisons against Forest and Low-productive soils.

A PICRUSt2 functional prediction analysis was then performed to assess differences in predicted microbial functions (Figure 6). PERMANOVA based on Bray–Curtis distances of PICRUSt2 unstratified pathway abundances showed no significant differences in overall functional composition among productivity levels (R^2^ = 0.12, p = 0.565) (Figure 6 - A). This indicates that the global functional structure of the soil microbiome is largely conserved across Forest, High-productive, and Low-productive sites. However, superpathway of demethylmenaquinol-6 biosynthesis II *(PWY-7373)* was significantly enriched in in high- and low-productive sites compared to Forest soils (MaAsLin2, q < 0.25) (Figure 6 - B). Furthermore, taxonomic contribution analysis revealed that genera such as *Sulfuricurvum, Paenibacillus, Cohnella*, and *Ammoniphilus* dominated the pathway in productive soils, while Forest sites showed less representation (Figure 6 - C).

**Figure 6.**
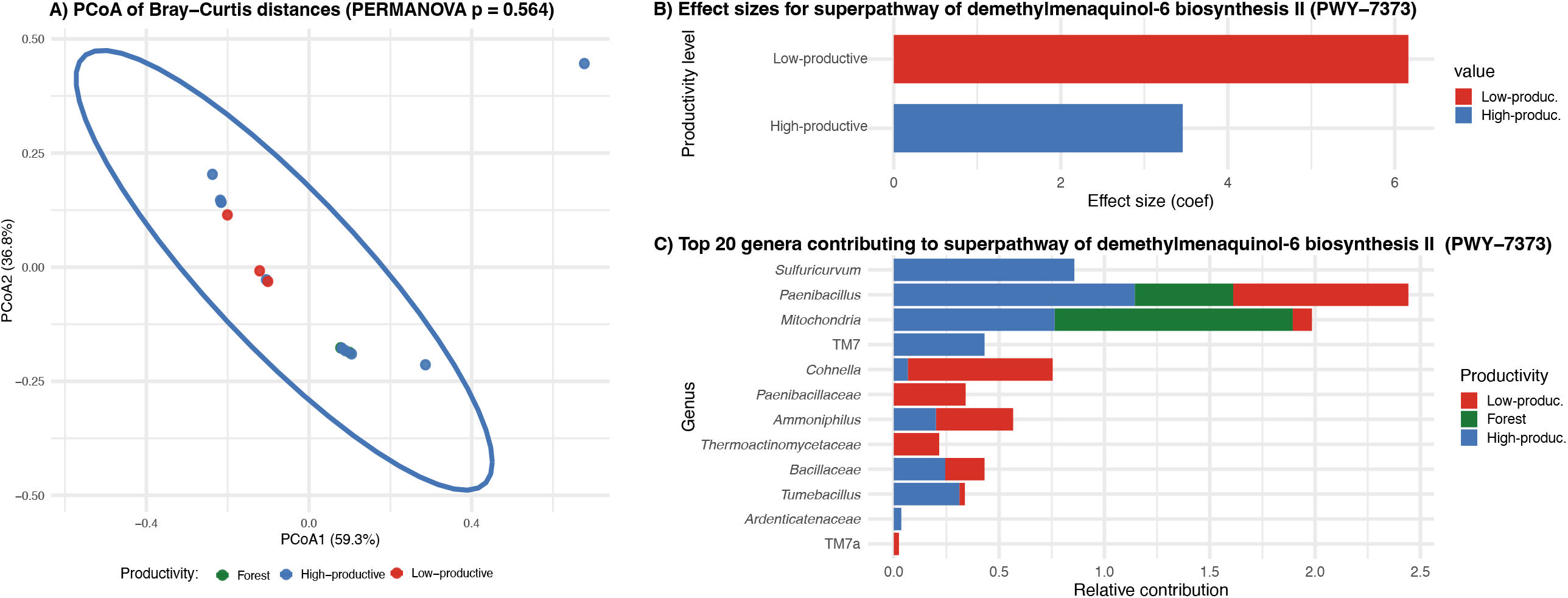
Principal coordinate analysis (PCoA) based on Bray–Curtis distances and differential abundance and taxonomic contributions of the superpathway of demethylmenaquinol-6 biosynthesis II (PWY-7373) across productivity levels. A) Principal coordinate analysis (PCoA) based on Bray–Curtis distances of predicted functional pathways generated using PICRUSt2. Samples showed substantial overlap among productivity groups, and PERMANOVA analysis indicated no significant differences in global functional profiles (R^2^ = 0.095, p = 0.194) (B) MaAsLin2 effect sizes showing enrichment of the pathway in Good and Bad sites relative to Forest. (C) Top 20 genera contributing to the pathway, colored by productivity level. The pathway is mainly driven by genera such as *Sulfuricurvum, Paenibacillus*, and *Cohnella* in low-productivity soils, indicating enhanced fermentative activity under degraded conditions.

Thus, when combining taxonomical profiling and functional prediction, these observations reflect that taxonomical changes are backed by high functional redundancy within the community.

## Discussion

The use of fertilizers has propelled agricultural output over the last decades; however, its use has also been associated to soil degradation and greenhouse gas emissions^34^. Therefore, the development of biotechnologies that allows crop to better utilize fertilizers has the potential to decarbonize agriculture while preserving productivity. Here, we performed a deeper evaluation of the predicted microbial functional potential of a previously developed microbiome-based fertilizer additive (“humus”), commercially applied in South America^22^. We also provide a first exploratory report of soil microbiomes from agricultural and forest sites in Paraguay.

Although PERMANOVA analysis did not detect significant differences in overall community composition between root- and humus-associated microbiomes, MaAsLin2 analysis identified the enrichment of specific bacterial lineages within roots, including members of the genus *Sphingomonas*, env.OPS_17 (family Sphingobacteriales), and Solirubrobacterales-related taxa. The enrichment of *Sphingomonas* is particularly noteworthy, as members of this genus are frequently associated with rhizosphere competence, stress tolerance, degradation of complex organic compounds, and plant growth promotion^35,36^. Similarly, taxa affiliated with Solirubrobacterales are commonly detected in soil and plant-associated ecosystems, including root endophytic environments^37^.

The humus functional profiling via PICRUSt2 prediction revealed enrichment in carbon degradation and central metabolism pathways, typical of compost-like systems ^4,38,39^. However, it also shared over 350 pathways with the root microbiome, underscoring their potential functional similarity. The enrichment of plant growth-promoting (PGP) pathways in root and humus, including auxin biosynthesis, siderophore production, and phosphate solubilization, mirrors similar findings with plant-associated PGPR consortia^25,40–43^. Their resemblance suggests that the humus might act as a functional analog of the rhizosphere microbiome, promoting nutrient cycling and plant development through microbial metabolism. Interestingly, a previously reported meta-analysis ^44^ highlighted that employing complex microbial consortia, over single strains, results in more consistent and effective improvements in soil fertility. From an ecological perspective, the effectiveness of complex microbial consortia is often attributed to complementary metabolic capabilities and partial functional redundancy among community members, reinforcing the biotechnological value of microbial diversity^45^.

Thus, this study provides the first detailed taxonomic and predicted functional characterization of a commercial microbiome-based fertilizer additive currently deployed across more than 1.4 million hectares in South America. Future work, including metagenomic and metatranscriptomic analysis, will focus on elucidating the exact microbial taxa and genes that might be involved in the results observed in the field, whereby the humus use leads to higher crop yields with lower fertilizer input.

In terms of the soils analyzed, although alpha diversity remained similar across productivity groups, beta diversity analyses revealed significant differences between high and low productive soils. The absence of significant differences in alpha diversity suggests that productivity was not associated with a generalized loss of microbial richness, but rather towards differences in the community structure. Furthermore, the differential abundance analysis indicated that 19 and 11 genera were significantly enriched in low productive and forest soils, respectively, and only one genus enriched in high productive soils. Forest soils were enriched in genera such as *Pseudomonas, Lysobacter, Cellvibrio, Terrabacter*, and *Hyphomicrobium*, many of which have previously been associated with rhizosphere interactions, organic matter turnover, and microbial competition in undisturbed or carbon-rich soils^46,47^. The enrichment of *Cellvibrio* in forest soils is consistent with previous reports linking this genus to cellulose degradation and turnover of plant-derived organic matter in soil ecosystems^48^. In contrast, low-productivity soils were enriched in taxa including *Bacillus, Paenibacillus, Lysinibacillus, Nitrosomonas, Methylocapsa*, and *Methylovirgula*. Several of these genera are commonly reported in nutrient-limited or environmentally stressed soils and are characterized by stress tolerance or specialized metabolic strategies^49,50^. For example, the enrichment of *Nitrosomonas* may indicate altered nitrogen cycling dynamics in low-productivity soils, whereas methanotrophic genera such as *Methylocapsa* and *Methylovirgula* are frequently associated with oligotrophic environments.

Despite these taxonomic differences, predicted functional profiles inferred using PICRUSt2 showed limited differentiation among groups, as beta diversity analyses were not significant. Similarly, MaAsLin2 identified only a single differentially abundant MetaCyc pathway (PWY-7373), which was enriched in agricultural soils. This corresponds to the superpathway of demethylmenaquinol-6 biosynthesis II, involved in the microbial synthesis of menaquinones (vitamin K2-related compounds) that function as electron carriers in bacterial respiratory chains. This predicted function may be associated with agricultural management conditions rather than productivity alone, as it plays an important role in anaerobic and facultative anaerobic respiration, particularly under fluctuating redox conditions^51^. Increased representation of this pathway in managed soils may therefore reflect shifts in microbial respiratory strategies associated with cultivation, fertilization, soil disturbance, or changes in oxygen and nutrient availability.

The apparent discrepancy between taxonomic and predicted functional variation is consistent with the concept of functional redundancy in soil microbiomes, where phylogenetically distinct taxa may encode overlapping ecological functions^52^. Similar patterns have been reported in other soil ecosystems, in which substantial compositional turnover occurs without major changes in predicted community function^53^. However, because PICRUSt2 predictions are inferred from marker gene data rather than directly measured through metagenomics or transcriptomics, these results should be interpreted as initial exploratory evaluation. Future shotgun metagenomic or transcriptomic analyses would help determine whether menaquinone biosynthesis is actively associated with agricultural soil management and productivity in these systems.

In conclusion, this work provides the first detailed taxonomic and predicted functional characterization of a commercial microbiome-based fertilizer additive currently used at large agricultural scales in South America. The observed overlap between humus and root-associated microbiomes along with the enrichment of plant-growth-promoting functions, suggests that this consortium may act as a rhizosphere-like microbial community capable of providing functions typically associated with plant-associated microbiomes. This provides a potential ecological framework for understanding the humus agronomic performance. Furthermore, by characterizing microbial communities across forest and agricultural soils in eastern Paraguay, this study establishes an initial baseline for future investigations of soil microbiome structure and function in one of South America’s most important agricultural regions. Future studies will include larger and longitudinal sampling and integrate multi-omics approaches to confirm functional expression and nutrient cycling dynamics.

## Acknowledgements

We would like to thank the Department of Biotechnology and the Faculty of Exact and Natural Sciences, National University of Asuncion (FACEN-UNA) for providing access to laboratory resources in molecular biology. The work was partially supported by the “Convenio de Formacion” between FACEN-UNA and MicroBios S.A. We would also like to thank Mr. Elton Tencaten for assisting in sample collection, and Mauricio Molinas, Lourdes Cardozo and Martin Nunez for sample processing. We would like to thank Bjoern Buss for proof-read assistance.

## Author contribution

WJSE conceived and designed the study, performed data analysis, and wrote the manuscript. WJSE also coordinated sample acquisition and laboratory work.

## Methods

This study aimed to characterize the taxonomic composition and predicted functional potential of a commercial microbiome-based fertilizer additive (Tiroleo Humus) and to compare microbial communities across agricultural and forest soils from eastern Paraguay (Department of Itapúa).

### Sample collection and gDNA extraction

#### Soil sampling

Soil samples were kindly provided by Tiroleo S.A. (Tiroleo, Paraguay) from their customers, as they routinely obtain for chemical and other microbiological analysis. These samples were collected in 50 mL sterile Falcon tubes, within the first 20 cm of soil’s top layer, and stored at -80°C until DNA extraction. For each plot, 50 samples separated 30 m from each other were collected and combined, representing one location. Their location was in the department of Itapúa in Paraguay and the fields were categorized as high- and low-productivity, referring to farmers reports regarding their crop yields. The remanent forest corresponds to areas adjacent to those commercial fields. The exact sample conditions and locations, including coordinates, are presented in Supplemental Material 3.

#### Humus sampling

Three samples were collected at the end of the three-week solid-state fermentation process from the surface, middle, and bottom sections of the solid-state fermentation boxes. Two root-associated microbiome samples previously analyzed in the original study were included for comparison.

DNA extraction was performed using the DNeasy Power Soil Pro Kit (Qiagen, Germany), following manufacturer’s instructions.

### 16S rRNA sequencing and analysis

#### Sequencing

Twenty ng/µL of DNA was used per sample. 16S rRNA gene sequencing was performed by Azenta Life Sciences (South Plainfield, NJ, USA) using their 16S-EZ service. Genomic DNA samples were subjected to proprietary multiplex PCR amplification targeting the V3 and V4 hypervariable regions of the bacterial 16S rRNA gene. Following amplification, a second PCR was performed to incorporate sample-specific barcodes and Illumina sequencing adapters. Libraries were pooled, quality-controlled, and sequenced on an Illumina platform using 2 × 250 bp paired-end chemistry. Because the amplification strategy relies on proprietary primer sets, primer sequences are not publicly available.

### Bioinformatics Analysis

Raw paired-end FASTQ files were processed in R using the DADA2 pipeline. Forward and reverse reads were quality filtered, denoised, merged, and screened for chimeric sequences following standard DADA2^55^ workflow, including merging paired ends, quality filtering, error correction, and chimera detection procedures. Amplicon sequence variants (ASVs) were assigned taxonomy using the SILVA v138 reference database. Alpha diversity with respect to Shannon index was estimated using R at a rarefaction depth of 5,000 sequences per subsample. Beta diversity estimates were calculated within R using Bray-Curtis dissimilarity between samples at a subsampling depth of 10,000. Samples containing fewer than 10,000 reads were excluded from beta diversity and downstream analyses to reduce potential biases associated with under-sampling and inflated sparsity. Principal coordinate analysis (PCoA) was used for ordination visualization. Group differences were assessed using PERMANOVA, and homogeneity of dispersion was evaluated using PERMDISP. Predicted microbial functional profiles were inferred using PICRUSt2 v2.6.3. ASVs were placed into a reference phylogeny and MetaCyc pathway abundances were inferred using the default PICRUSt2 workflow. Results were summarized and visualized through principal coordinate analysis, and significance was estimated as implemented in R. We used MaAsLin2 (Microbiome Multivariable Association with Linear Models) to identify differentially abundant taxa and pathways. Separate models were fitted for comparisons based on Type (Root vs Humus, for the humus section) and Productivity (Forest, High-productivity, and Low-productivity soils, for the soils section). High productive and forest soils were set as reference level for taxonomic and predicted functional, respectively. Statistical significance was defined as q < 0.25. Taxonomic abundance data were normalized using total sum scaling (TSS) and log-transformed prior to association testing, and with a minimum of 10% prevalence. Features with q-values < 0.25 and effect sizes (coef) indicating higher or lower abundance were retained for comparison.

#### A.I. Disclosure

Large Language Model (LLM) was used to assist with English grammar or for rewriting paragraphs to lower the words count. All scientific interpretations, analyses, and conclusions were generated and reviewed by the author.

## Data availability statement

Sequence data have been deposited in the NCBI Sequence Read Archive (SRA) under BioProject accession number PRJNA1305027.

Online at https://dataview.ncbi.nlm.nih.gov/object/PRJNA1305027?reviewer=k16f9mso524liksqqrci10sc0s until release, after which the link will be: https://submit.ncbi.nlm.nih.gov/subs/sra/SUB15531780/overview, accession number PRJNA1305027.

